# An immunoediting map of human cancers

**DOI:** 10.1101/2024.06.08.598035

**Authors:** Rui Gan, Xianwen Ren

## Abstract

Understanding how cancer immunoediting sculpts tumor microenvironments is essential to disentangling tumor immune evasion mechanisms and developing immunotherapies. Here, we construct a comprehensive immunoediting map of human cancers via single-cell deconvolution of 11057 tumor-derived samples across 33 cancer types from TCGA and comparison with 17382 healthy samples across 30 tissues from GTEx. The map covers >1000 different cell states across all the major immune cell types. Mast cells, megakaryocytes, macrophages, neutrophils, plasma cells and T cells are up-regulated across a wide range of tumor types while natural killer cells and platelets are down-regulated in most tumor types, suggesting common cancer immunoediting events. While tumor heterogeneity is higher than the normal corresponding tissues, significant immune homogeneity exists among different tumor types compared with the distinct immune composition among normal tissues and organs. Our study provides a new holistic perspective to understanding cancer immunoediting. Our findings may provide important hints for developing novel cancer immunotherapies, and the high-resolution immunoediting map may serve as a rich resource for further pan-cancer investigation.

## Introduction

Cancer is a major cause of death worldwide. According to the latest data from the World Health Organization (WHO), cancer leads to about 10 million deaths, accounting for about 17% deaths in 2020. Human immune system plays a vital role in protecting the body against cancer (*1*). However, cancer immunoediting frequently occurs, leading to the acquisition of edited immunogenicity and immune escape by the tumors (*2*). During this process, carcinogenesis can trigger numerous functional alterations and compositional shifts within the immune system (*3*). Malignant cells are able to evolve different mechanisms to escape the host immune pressure, driving tumor progression and therapeutic resistance (*4*). While single-cell analysis of individual tumor types and pan-cancer analysis yield profound insights into tumor microenvironments and the potential cancer immunoediting mechanisms, healthy controls were generally set as tumor-adjacent tissues rather than true healthy tissues and organs (*5–7*). It is still missing of a systematic comparison between tumors and true healthy tissues. In this study, we elaborate a single-cell deconvolution strategy to investigate how cancer immunoediting alters the immune composition in the tumors relative to true healthy tissues. By single-cell deconvolution of 17382 RNA-seq samples from The Genotype-Tissue Expression (GTEx) using Redeconve as the algorithmic workforce and Tabula Sapiens as the single-cell reference, we have constructed an immune map of human body and named it as RedeImmunoMap (*8–11*). RedeImmunoMap not only demonstrates high agreement with the measured immune cell densities from literatures based on histology and flow cytometry techniques (*12*), but also provides unprecedented resolution to investigate the cellular composition of human normal tissues and organs across tens of thousands of samples. With RedeImmunoMap as a healthy control, here we perform single-cell deconvolution analysis based on the 11057 tumor samples across 33 cancer types from The Cancer Genome Atlas (TCGA) project to investigate the common and different cancer immunoediting strategies across different tumor types (*13*). The results deepen our understanding of the immunoediting landscape during carcinogenesis, facilitating the development of innovative therapeutic immunomodulation strategies.

## Results

### Overview of the immunoediting map of human cancers

In our previous work, we constructed RedeImmunoMap, a comprehensive human immune map by single-cell deconvolution of 17382 GTEx samples across 30 major tissues with Redeconve using sampled 5002 cell states from Tabula Sapiens. We have validated our results on the literation data. Based on these efforts, we performed sing-cell deconvolution analysis on 11057 TCGA samples across 33 cancer types using the same single-cell reference as RedeImmunoMap (**Fig. 1**). After filtering adjacent, metastatic or specific-type samples, we obtained 9875 primary tumor samples, in which 1144 immune and 223 endothelial cell states were detected, including 126 B cells, 93 plasma cells, 325 T cells, 28 natural killer (NK) cells, 24 natural killer T (NKT) cells, 367 monocytes/macrophages, 33 dendritic cells (DCs), 99 neutrophils, 3 eosinophils, 3 basophils, 33 mast cells, 9 innate lymphoid cells (ILCs) (**Fig. S1**). By integrating with RedeImmunoMap, we constructed a comprehensive map of cancer immunoediting by dissecting the immune perturbations in 8747 tumor samples across 26 cancer types and 12141 healthy samples from 20 paired normal tissues, involving with 1232 immune and 259 endothelial cell states (**Fig. 1**). We further evaluated the performance of deconvolution results by calculating cosine similarities between the true and reconstructed gene expression profiles. The averaged similarity scores exceeded 0.9 for most tissues, proving the high quality of deconvoluted abundance. We also observed widespread significant decrease of similarity scores between tumor and corresponding healthy samples in most tissues due to carcinogenesis (**Fig. 2A**). Overall, we successfully reconstructed the whole immune and endothelial cellular compositions of all major immune cell types in the tumor and healthy samples, providing a comprehensive resource to systematically investigate the cancer immunoediting process across various cancer types (**Fig. 2B**).

**Fig. 1.**
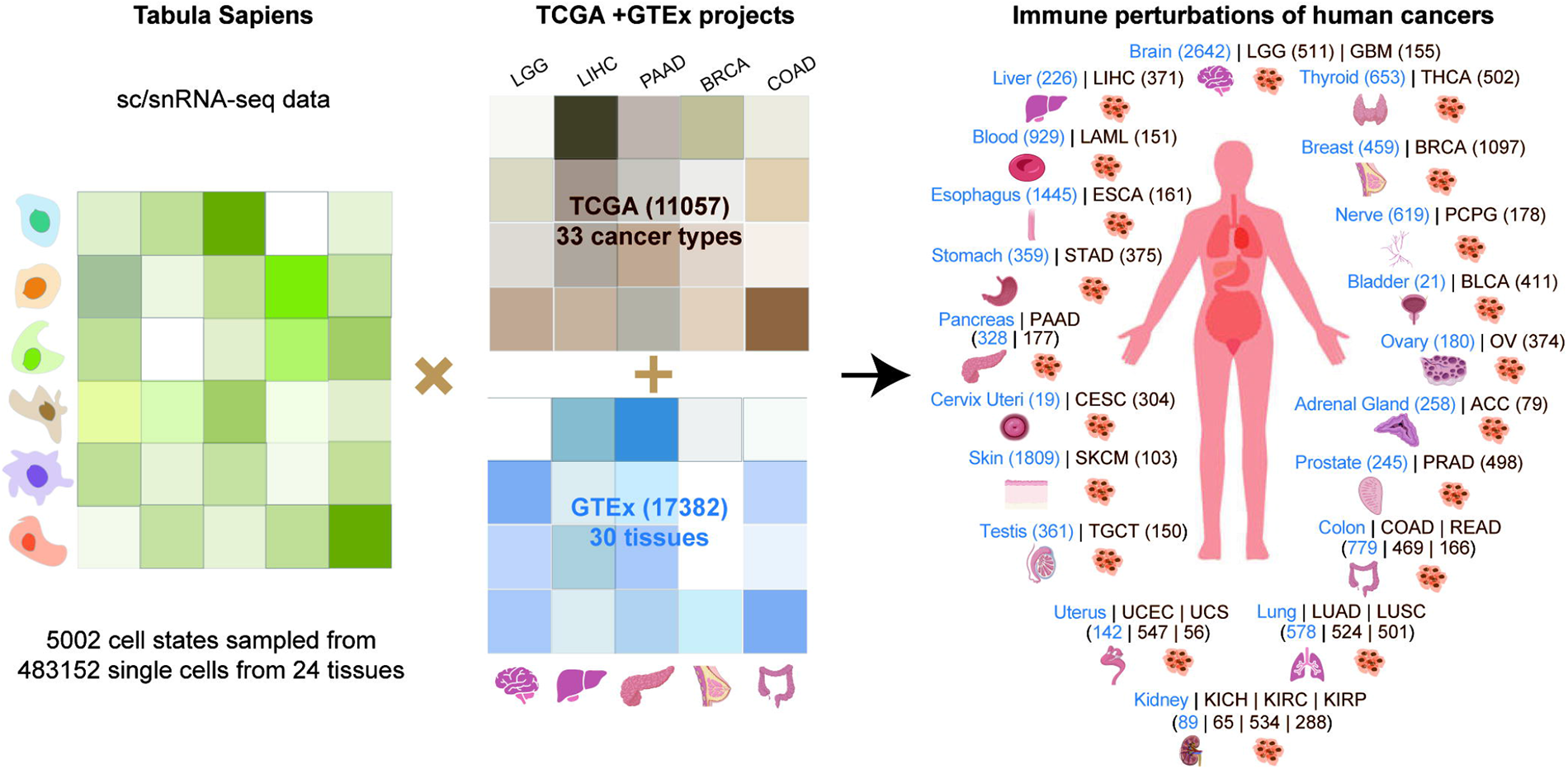
Overview of the immunoediting map. The scRNA-seq data from Tabula Sapiens and bulk RNA-seq data from GTEx and TCGA were integrated by Redeconve. The estimated cell abundance matrix was analyzed to depict the immune abundance perturbations between 20 healthy tissues and 26 cancer tissues. The tissue figures were quote from Gan et al. (*8*).

**Fig. 2.**
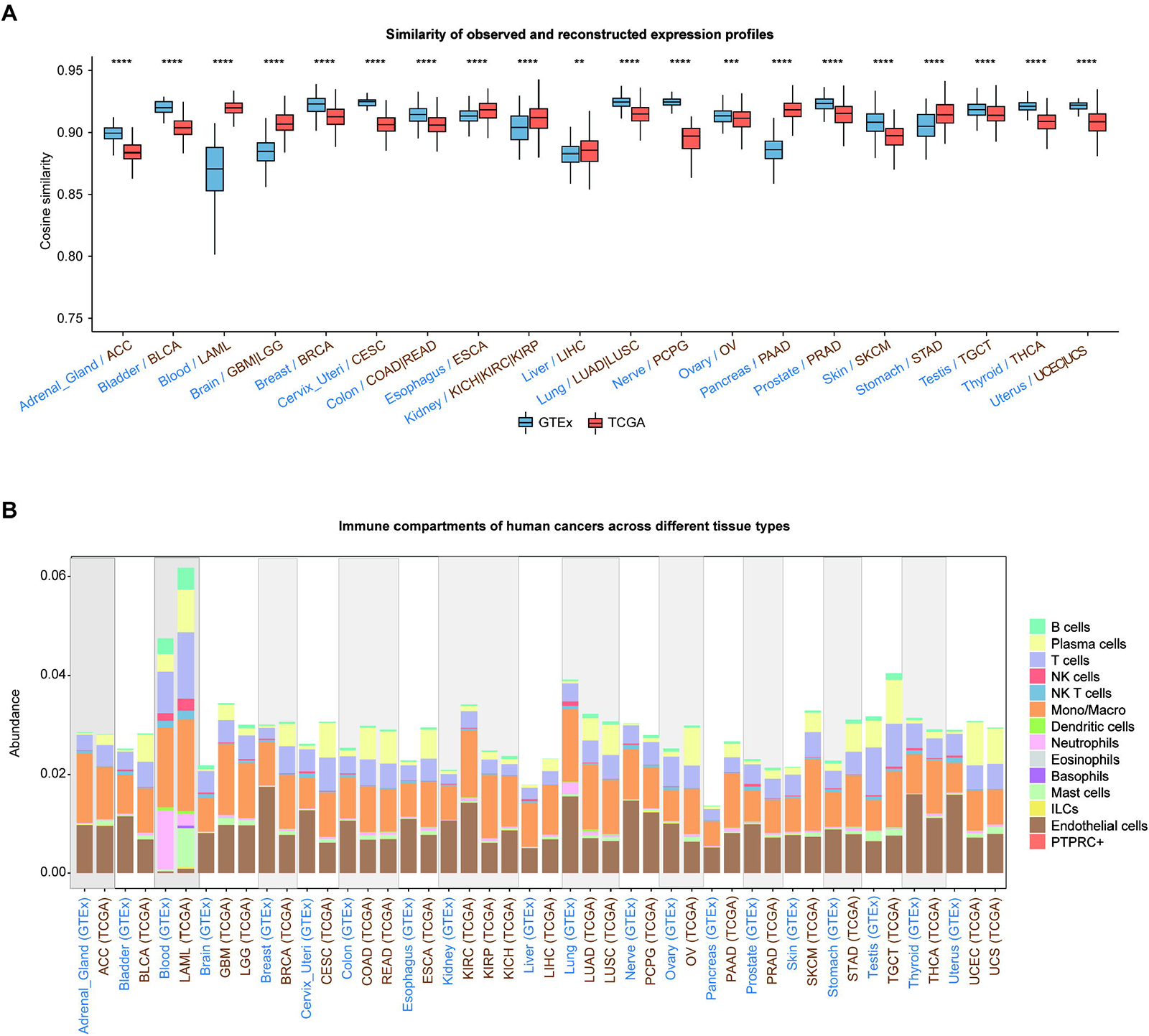
Performance of single-cell deconvolution. A, Cosine similarities of tissue samples between observed and reconstructed expression profiles in 20 healthy tissues and 26 cancer tissues (Wilcoxon rank-sum test). B, Bar plot showing the cellular compartments of each tissue. The abundance of each cell state was averaged for each tissue. \**P* < 0.05, \*\**P* < 0.01, \*\*\**P* < 0.001, \*\*\*\**P* < 0.0001.

### Cancer-induced mast cell expansion across diverse tumor types

Mast cells play a vital role in regulating both innate and adaptive immune responses by secreting a wide array of biologically active factors (*14*). Previous studies indicate that mast cells can promote the impairment of tumor progression by secretion of pro-inflammatory mediators or reversely lead to tumor progression by modulating the immune responses (*15, 16*). However, how mast cell perturb the tumor microenvironment have not been systematically investigated across different cancer types. Here, we thoroughly investigated mast cell compositional changes from healthy tissues to tumors across 26 cancer types (**Fig. 3**). We observed that mast cell abundance significantly increased in most cancer types compared with the healthy samples except testicular cancer (**Fig. 3A**). Furthermore, we investigated the distribution of mast cell perturbation in the males and females across diverse cancer types. Consistent expansion of mast cells is observed for both sexes during carcinogenesis (**Fig. 3B**). Additionally, we characterized the mast cell abundance alterations across various cancer types in different age groups, showing a strong evidence of mast cell expansion for all ages (**Fig. 3C**). Together, our results revealed that mast cells universally increased induced by tumor burden for almost all cancer types, potentially serving as a novel immunotherapeutic target in cancer treatment.

**Fig. 3.**
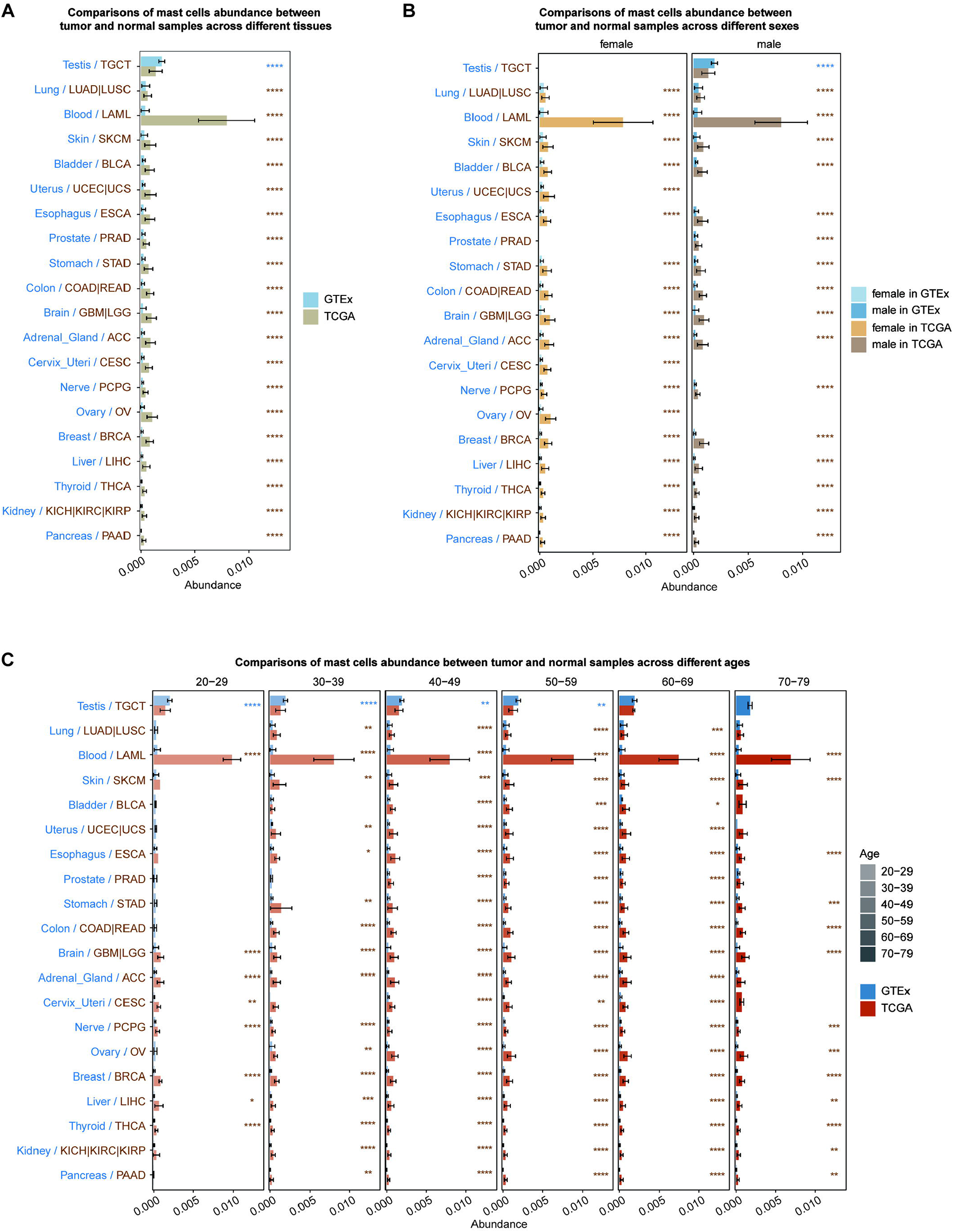
Distribution of cancer-induced mast cell perturbation across different tissues, sexes and ages. A, Bar plot showing the distribution of cancer-induced mast cell perturbation between 20 tissues and 26 cancer tissues ranked by their averaged mast cell abundance in the healthy samples. Error bars were plotted by mean±SD for A, B and C. The mast cell abundance of each RNA-seq sample was aggregated for all mast cell states for A, B and C. B, Distribution of mast cell perturbations in the males and females. C, Distribution of mast cell perturbations across different age groups. Wilcoxon rank-sum tests were performed for cell type abundance between healthy and tumor samples for each tissue. \**P* < 0.05, \*\**P* < 0.01, \*\*\**P* < 0.001, \*\*\*\**P* < 0.0001.

### Depletion of natural killer cells in a wide range of human cancers

NK cells are effector lymphocytes mediating innate immune response with natural cytotoxicity and cytokine-producing effector functions (*17*). NK cells can directly recognize and kill tumor cells as well as mediate anti-tumor immunity of other immune cells (*18*). For example, Glasner et al. demonstrated that the activation of the NK natural cytotoxic receptor 1(Ncr1) (mouse) and NKp46 (human) modulated fibronectin 1 expression on tumor cells by triggering IFNγ production, consequently preventing tumor metastasis (*19*). Several studies have observed decreased expression of activating receptors in peripheral NK cells from patients with breast cancer, NSCLC, neuroblastoma or gastrointestinal stromal tumors, compared with healthy samples, which revealed the impaired cytotoxicity and specificity of NK cells in the tumor microenvironment (*20–24*). Therefore, we explored the NK cell perturbations in the tumor microenvironment across 26 cancer types in different sex and age groups (**Fig. 4**). Our results indicated that almost all cancers exhibited a significant depletion of NK cells, except for blood, testis, kidney and pancreas (**Fig. 4A**). Notably, we found the NK cell perturbations exhibited consistent tendency between males and females in almost all cancer types except esophageal carcinoma (ESCA) (**Fig. 4B**). We further found that older ESCA patients exhibited reduction of NK cells, compared with the younger patients (**Fig. 4C**). In addition, similar phenomena were observed in brain, breast and skin tumors, revealing the detrimental impact of aging on anti-tumor capacity of NK cells. Overall, our results point a potential strategy of NK cell-based immunotherapy in most cancer types.

**Fig. 4.**
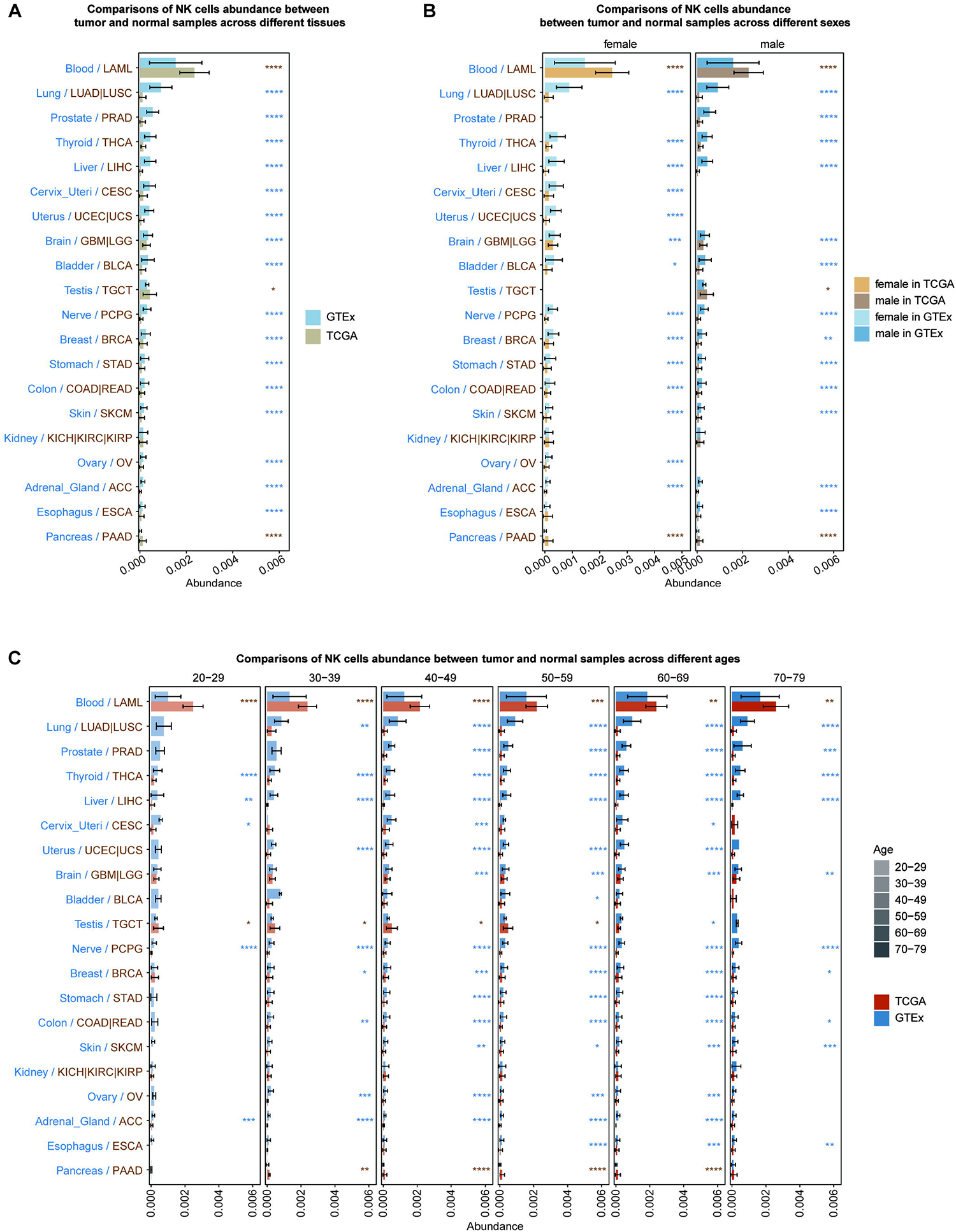
Distribution of cancer-induced NK cell perturbation across different tissues, sexes and ages. A, Bar plot showing the distribution of cancer-induced NK cell perturbation between 20 tissues and 26 cancer tissues ranked by their averaged NK cell abundance in the healthy samples. Error bars were plotted by mean±SD for A, B and C. The NK cell abundance of each RNA-seq sample was aggregated for all NK cell states for A, B and C. B, Distribution of NK cell perturbations in the males and females. C, Distribution of NK cell perturbations across different age groups. Wilcoxon rank-sum tests were performed for cell type abundance between healthy and tumor samples for each tissue. \**P* < 0.05, \*\**P* < 0.01, \*\*\**P* < 0.001, \*\*\*\**P* < 0.0001.

### Cancer immunoediting of other immune cell types across different tumor types

We further investigated the cancer-induced perturbations of all immune cell types plus endothelial cells across diverse tissues and cancer types beyond mast cells and NK cells (**Fig. S2 to S20**). Firstly, a significant increase of all immune cell abundance was observed in tumor samples, compared with the healthy samples across almost all tissues, showing the recruitment of immune cells in the tumor microenvironment (**Fig. S2**). We further found plasma cells, T cells, CD4^+^ T helper cells, CD8^+^ T cells, regulatory T cells, monocytes/macrophages and neutrophils and basophils plus mast cells exhibited significant induction while NK cells, NKT cells and mucosal associated invariant T (MAIT) cells showed a widespread reduction in different sex and age groups in the tumor microenvironment, compared with the healthy samples. In addition, B cells, γδ T cells, dentritic cells and ILCs exhibited tumor-specific perturbations (**Fig. S2 to S20**). For B cells, we observed significant depletion in brain, adrenal gland, ovary, cervix uteri and uterus tumors as well as stronger B cell perturbations in the brain tumors of males and breast tumors of females (**Fig. S3**). Plasma cells exhibited a global induction in all cancer tissues, revealing the potential plasma cell clonal expansion in the tumor microenvironment (**Fig. S4**). We also observed T cells decreased in testis, ovary and thyroid tumors (**Fig. S5**). It was interesting that CD4^+^ T helper cells sharply decreased while CD8^+^ T cells substantially increased in the testicular germ cell tumors (TGCT) and ovarian serous cystadenocarcinoma (OV) (**Fig. S6 and S7**). MAIT cells showed a sharp reduction in testis, ovary and brain tumors but significant induction in acute myeloid leukemia (LAML) and pheochromocytoma and paraganglioma (PCPG) (**Fig. S10**). Monocytes/macrophages as the largest population of immune cells were enriched in most tumor types except for lung, adrenal gland and nerve tumors while DCs mainly diminished in ovary, skin, adrenal gland and nerve tumors (**Fig. S12 and S13**). Moreover, we found neutrophils drastically expanded in most tumor types except for blood and lung after tumor infiltering (**Fig. S14**). Comparatively, basophils and ILCs sharply increased in leukemia patients (**Fig. S16 and S17**). Additionally, we performed comparisons of PTPRC^+^ITGA2B^+^ and PPFB^+^PF4^+^ cell abundance between tumor and healthy samples, respectively representing megakaryocytes and platelets (*25*) (**Fig. S18 and S19**). We observed global induction of PTPRC^+^ITGA2B^+^ cells but extensive reduction of PPFB^+^PF4^+^ cells, revealing that the abnormal megakaryocytes failed to functionally generate platelets in the tumor microenvironment. Furthermore, we found endothelial cell abundance significantly decreased in most tumor tissues (**Fig. S20**). Overall, we comprehensively delineated the perturbation patterns of diverse immune cell types in various cancer types induced by immunoediting, as well as their correlations with sex and age, aiming to guide the development of targeted immunotherapies.

### Immune homogeneity of different cancer types in contrast to the immune heterogeneity of different normal organs

We further comprehensively depicted the associations and differences of immune plus endothelial states between different tumor and healthy tissues. The correlation analysis revealed a significant difference of immune features between the tumors and healthy tissues, indicating the corrupted states of immune organization in the tumors (**Fig. S21**). Notably, we observed high correlations between cancer types with similar histopathologic features or from relevant tissue sites. For examples, brain lower grade glioma (LGG) and glioblastoma multiforme (GBM) exhibited a correlation of 0.84. Testicular germ cell tumors (TGCT), ovarian serous cystadenocarcinoma (OV), uterine corpus endometrial carcinoma (UCEC), uterine carcinosarcoma (UCS) and cervical squamous cell carcinoma and endocervical adenocarcinoma (CESC) highly correlated in immune compositions. In addition, high correlations of immune compositions were also observed between adrenocortical carcinoma (ACC) and kidney cancers (KIRP, KICH and KIRC), liver hepatocellular carcinoma (LIHC) and cholangiocarcinoma (CHOL) as well as among gastrointestinal cancers including rectum adenocarcinoma (READ), colon adenocarcinoma (COAD), stomach adenocarcinoma (STAD), esophageal carcinoma (ESCA) and pancreatic adenocarcinoma (PAAD). Altogether, we revealed the immune associations between different cancer types, which were potentially related with histopathologic features.

Previous studies have reviewed that tumors will drive extensive disruption of hematopoiesis, exhibiting prominent immune cell perturbations, thereby leading to local immunosuppression (*26–28*). Here, we globally demonstrated the overall immune distributions across diverse healthy tissues and tumor types, exhibiting great changes of immune organization (**Fig. 5A**). We visualized the immune features of healthy and cancer tissues into the same two-dimensional space using t-distributed stochastic neighbor embedding (t-SNE) (**Fig. 5B**). The t-SNE plots revealed the strong immune dysregulation in tumor tissues in cancer evolution. Each tissue module exhibited specific and organized immune features in healthy samples, playing steady functions in immune response. Comparatively, most tumor types were disordered, undergoing dramatic compositional changes during tumor progression. For examples, acute myeloid leukemia (LAML) exhibited severe immune corruption with an extremely low correlation of 0.26, compared with healthy blood samples (**Fig. S21**). The female reproductive organs including ovary, uterus and cervix uteri exhibited unsolid immune organizations during tumor progression, showing low correlations of 0.33, 0.34 and 0.36 with the corresponding tumor tissues (**Fig. 5C**). Comparatively, we discovered that a minority of tissues exhibited minor alterations in the immune composition in the progress of cancerization. For examples, adrenal gland and ACC, liver and LIHC as well as brain and brain tumors including LGG and GBM exhibited similar immune features with moderate correlations of 0.77, 0.74 and 0.72, indicating that these tissues harbored relatively stable immune organization and homeostasis (**Fig. 5D**). In summary, our results demonstrate higher immune homogeneity of different cancer types in contrast to the immune heterogeneity of different normal organs.

**Fig. 5.**
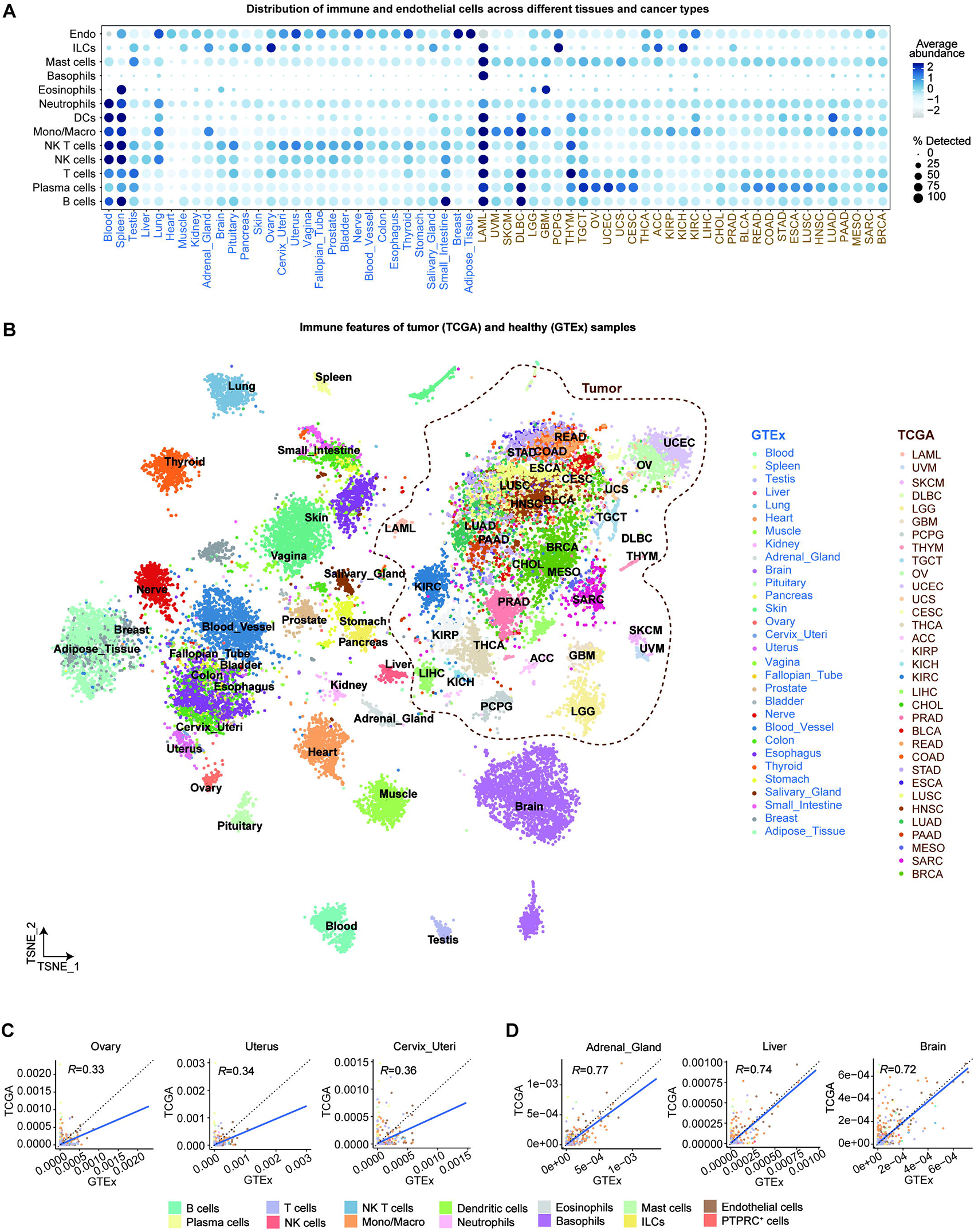
Immune associations between human tissues. A, Dot plot showing tissue-specific abundant cell types colored by scaled mean abundance across 30 healthy tissues and 33 cancer tissues for each major cell type. The size of points indicted the percentage of samples detecting the major cell type. B, t-SNE visualization of 17382 healthy samples and 9875 tumor samples across 30 healthy tissues and 33 cancer tissues based on their cell state abundance.

### Higher inter-individual immune heterogeneity for different cancer types compared with healthy tissues

The human immune system exhibits significant variation among individuals, yet it remains relatively stable over time for each individual person. It has been reported that the human immune system variation serves as a key factor influencing variations in the success of immunological therapies for cancer treatment (*29*). However, how immunoediting alters the inter-individual immune variation has not been reported yet. Therefore, we performed comprehensive comparisons for inter-individual variations of all immune plus endothelial cell states, measured by standard deviation of cell state abundance, between the healthy and corresponding tumor tissues (**Fig. 6**). Our results indicated that all the tumor tissues harbored significantly higher inter-individual immune heterogeneities except for LAML, compared with the healthy tissues (**Fig. 6A**). Pancreas and testis tumors exhibited excessive inter-individual immune heterogeneities than the normal tissues for almost all cell states. In addition, the level of immune variation observed in kidney, uterus, bladder and cervix uteri tumor tissues also significantly surpassed the corresponding normal tissues, reaching over 1.5 times higher (**Fig. 6B**). Altogether, our analysis revealed extremely high inter-individual immune heterogeneities among different cancer patients, uncovering the complicated process of immunoediting.

**Fig. 6.**
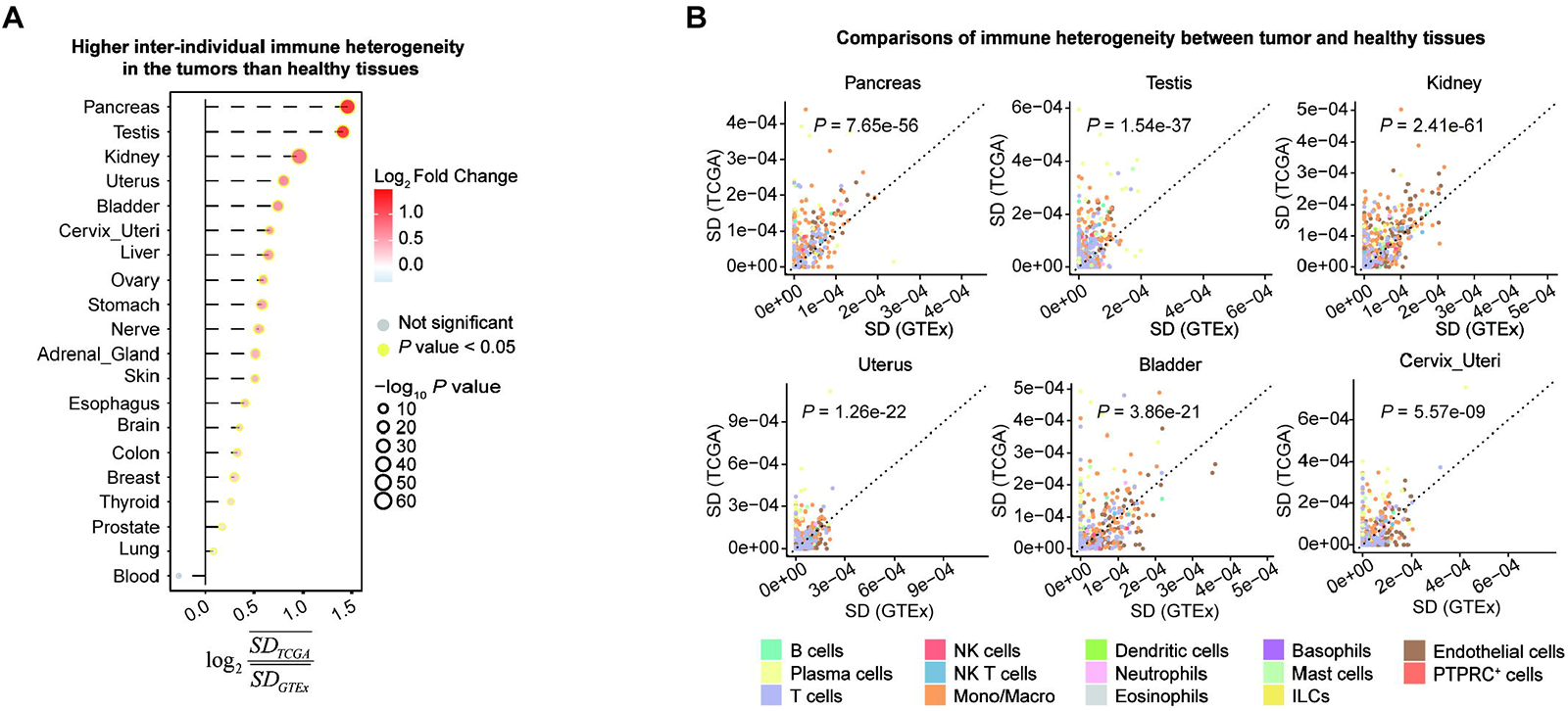
Comparisons of inter-individual immune variations between tumor and healthy tissues. A, Bubble plot showing the comparisons of standard deviation values for 1232 immune and 259 endothelial cell state abundance between TCGA and GTEx tumor samples by paired Wilcoxon rank-sum tests. The point size indicated the significance level and border color indicated whether *P* < 0.05. B, Scatter plots showing the standard deviation values for 1232 immune and 259 endothelial cell state abundance in TCGA and GTEx samples from pancreas, testis, kidney, uterus, bladder and cervix uteri. The dashed diagonal line represents y=x. The labeled *P* values represent the significance level of paired Wilcoxon rank-sum test.

## Discussion

Effective immunotherapeutic approaches require a complete understanding of cancer immunoediting process across different cancer types and populations. Therefore, it is essential to fully investigate the cancer-induced immune perturbations in order to develop novel immunotherapies for maintaining the balance of immune homeostasis and increase immune response against cancers. We previously have proved the success of integration of scRNA-seq and bulk RNA-seq datasets for constructing a comprehensive immune map of human body. In this study, we continued utilizing single-cell deconvolution methods of depicting immune organizations in diverse tumor types. By combining with previous immune map of healthy human bodies, we comprehensively depicted the immune perturbations in the immunoediting process across a wide range of cancer types in various sex and age groups. We found that the overall immune cell abundance significantly increased in the tumor microenvironment compared with the healthy tissues in the males and females. We further investigated the perturbation patterns of each immune cell type and the impact of sex and age on the perturbations. We found that plasma cells, T cells, CD4^+^ T cells, CD8^+^ T cells, monocytes/macrophages, neutrophils and mast cells exhibited significant enhanced enrichment within tumors across a wide range of tumor types while NK, NKT and MAIT cells showed a widespread reduction in almost all cancer types. Additionally, B cells, γδ T cells, dentritic cells and ILCs exhibited tumor-specific perturbation patterns across diverse tumor types and are influenced by sexes and ages. In addition, we found the perturbation pattern exhibited difference in various ages for particular tissues and cell types. For example, we observed that NK cell perturbation exhibited a correlation with age in esophagus, brain, breast and skin tumors, revealing the negative impact of aging on anti-tumor immunity. Furthermore, we depicted dramatic difference between healthy tissues and cancer tissues as well as the associations of different tumor types, probably related with histopathologic features of tumors. We also indicted the immune dysregulation in the tumor microenvironment but relative immune homeostasis in adrenal gland, liver and brain. We finally revealed the higher immune heterogeneity in the tumor microenvironment compared with the healthy tissues, partially explaining the great difficulties of immunotherapies. Our results provide a comprehensive landscape of immune response against cancer across diverse cancer types, of which the detailed alterations can be further investigated by means of *in silico* flowcytometry for example. The mechanisms driving such perturbations we reported remain unknown. Hence, future research will necessitate a rigorous exploration of the fundamental mechanisms underlying immunoediting process, aiming to precisely design therapeutic strategies capable of restoring a dysregulated immune system to a state of healthy immune homeostasis.

## Methods

### Public datasets

The bulk RNA-seq expression profiles and corresponding metadata information were downloaded from https://xenabrowser.net/datapages, including 11057 TCGA samples across 33 cancer types. The scRNA-seq reference originated from the Tabula Sapiens and deconvolution results of 17382 healthy RNA-seq samples were downloaded from https://zenodo.org/records/11057781, including 5002 cell states.

### Deconvolution of RNA-seq data

The bulk RNA-seq expression values were estimated by the Transcripts Per Million (TPM) measure, then log_2_ transformed. Further, the log-transformed bulk RNA-seq expression matrix and scRNA-seq expression matrix were together inputted into Redeconve to estimate the abundance of each cell state in each RNA-seq sample. The obtained abundance matrix was used for subsequent analysis.

### Comparative analysis of cell type abundance between healthy and cancer tissues

The cell type abundance of each RNA-seq sample was calculated by aggregation of corresponding cell state abundance. For each tissue, we performed Wilcoxon rank-sum tests between healthy samples and corresponding tumor samples. The tumor samples derived from primary solid or blood tumors filtered by sample name. The detailed information can be viewed from https://gdc.cancer.gov/resources-tcga-users/tcga-code-tables/sample-type-codes.

### Immune correlation analysis across tissues

The RNA-seq samples were visualized in two dimensions using t-distributed stochastic neighbor embedding (t-SNE) based on their abundance of 1232 immune and 259 cell states. Furthermore, the abundance of immune and endothelial cell states in each tissue was averaged and then used for pearson correlation analysis. We visualized the correlations between 30 healthy tissues and 33 cancer tissues via R package “corrgram” (*30*).

### Immune heterogeneity analysis across tissues

For each tissue, we calculated the standard deviation value among the corresponding samples for each cell state, which we used to measure inter-individual immune heterogeneity. We performed paired Wilcoxon rank-sum tests for cell state abundance between TCGA tumor and GTEx samples in each tissue.

## Acknowledgements

This work was supported by the National Natural Science Foundation of China (32022016 to X.R., 92159305 to X.R., and 31991171 to X.R.), National Key R&D Program of China (2022YFC3400904 to X.R.), and Changping Laboratory.

## Author contributions

X.R conceives and supervises this project. R.G. conducts the deconvolution and subsequent analysis of different cancers regarding the immune composition and associations with sex and age. R.G. and X.R. wrote the manuscript together.

## Data availability

The codes and corresponding data for Redeconve analysis as well as estimated cell abundance matrix were deposited at https://zenodo.org/records/11227690.

